# Cardiac-immune microniches programme macrophage states in the regenerating heart

**DOI:** 10.64898/2026.03.05.709830

**Authors:** Ehsan Razaghi, Selin Tüzüner, Trishalee Gungoosingh, Kerem Çil, Wan Shern Wong, Rita Alonaizan, Rebecca Richardson, Filipa C. Simões

**Affiliations:** Institute of Developmental and Regenerative Medicine, Department of Physiology, Anatomy and Genetics, University of Oxford; School of Physiology, Pharmacology and Neuroscience, University of Bristol

**Keywords:** Regeneration, Heart, Microniche, Macrophage, Microenvironment, Spatial transcriptomics, Zebrafish

## Abstract

Adult zebrafish regenerate their hearts after injury, a process that requires macrophages, yet how local tissue microenvironments instruct macrophage states and function remains unclear. Here we combine single cell RNA sequencing with Visium and high-resolution MERFISH spatial transcriptomics to map the cardio-immune landscape of homeostatic and regenerating zebrafish hearts. We identify a mpeg1.1+ compartment comprising macrophages, dendritic, B and NK-like cells, and show that injury establishes a macrophage-centred immune environment with transcriptional programmes spanning resident surveillance, damage sensing, inflammation, antigen presentation, resolution and metabolic support. Communication-aware spatial modelling reveals that these states are not randomly distributed but organised into discrete structural-immune microniches across the injury region, each defined by stereotyped cellular compositions and ligand-receptor circuits. Within a fibroblast-macrophage microniche, we uncover an il34-csf1ra-egr1 axis in which col12a1a+ il34+ fibroblasts promote an egr1 pro-regenerative macrophage state that couples fibrosis, vascular integrity and epicardial signalling. We show that disruption of this axis by csf1ra loss of function reduces macrophage-, endothelial- and epicardial-rich microniches, amplifying fibroblast-driven domains that shift macrophages towards stress and long-sustained inflammatory programmes, thereby biasing early injury response towards a pro-fibrotic state. Our work establishes spatially defined cardio-immune microniches as key organisers of macrophage function and regenerative outcome, providing a mechanistic framework and actionable targets for reprogramming cardiac repair.

## INTRODUCTION

Myocardial infarction triggers a rapid immune response that is essential for cardiac repair but frequently locks the adult mammalian heart into permanent scarring and progressive heart failure (1). In these settings, immune cells clear dead tissue and stabilise the ventricle, yet prolonged or dysregulated inflammation drives fibroblast activation, extracellular matrix deposition and irreversible loss of contractile myocardium (2, 3). In contrast, zebrafish, axolotls and other regenerative vertebrates mount an immune response that not only repairs the acute lesion but supports restoration of cardiovascular tissue and function through a tightly modulated immune response that favours cardiac regeneration (4–8). Macrophages emerge as central effectors of this regenerative response. Comparative work in zebrafish and medaka shows that even modest delays in macrophage recruitment are sufficient to impair cardiomyocyte proliferation, revascularisation and scar resolution (5). In salamanders, macrophage depletion causes premature fibroblast activation and formation of highly cross-linked, non-resolving scars (6). In zebrafish, resident cardiac macrophages with primitive origins expand after injury and regulate oxidative stress and extracellular matrix remodelling, which infiltrating monocyte-derived macrophages cannot replace (9, 10). Recruited macrophage subsets persist beyond the inflammatory phase and coordinate matrix degradation and tissue remodelling, for example via Mmp14 and Timp family proteases (11). These studies establish the functional importance of distinct macrophage populations as to their presence in response to injury, when they act and how they integrate local signals, as key determinants of whether hearts regenerate or permanently scar.

Macrophage behaviour is highly sensitive to the local environment, as shown by adoptive transfer experiments in zebrafish and mice where macrophages from regenerative settings promote fibrosis when placed into scar-prone environments and, conversely, adopt reparative programmes when moved into pro-regenerative niches (8). These findings argue that macrophage identity and function are not fixed, cell-intrinsic traits but rather emergent properties of signals received within specific tissue microenvironments.

The concept of such microenvironments, or niches, is now being refined by high-dimensional omics. In the human heart, integrated single cell and spatial transcriptomics have revealed region-specific cardiac niches, including epicardial immune niches enriched for plasma cells and conduction system niches with specialised glial and immune partners (12, 13). In non-regenerative mouse and human myocardial injury, spatial multi-omics has uncovered complex cardio-immune niches in infarcted and border zones, including Trem2-high macrophage-fibroblast circuits that restrain fibroblast proliferation and limit fibrosis (3, 14–16). Beyond the heart, immune microniches organising Treg function in the gut and Th2 differentiation in the lung have been mapped, showing that spatially restricted interactions between dendritic cells, macrophages and T cells are critical for tolerance and allergic inflammation (17–19). These studies shift the focus from individual cell types to niche-level cellular crosstalk, in which spatially confined cellular communities, their signalling circuits and their collective functions determine tissue-level responses.

Despite this progress, regenerative cardiac niches remain poorly defined at cellular and molecular resolution. In the zebrafish heart, single cell transcriptomics and imaging studies have catalogued diverse resident macrophage, dendritic and mononuclear phagocyte populations and established their importance for regeneration (9, 10, 20–23). Yet, we still lack a spatially resolved view of how these immune populations are organised relative to cardiac structural cells during regeneration and how niche-specific cellular circuits might instruct macrophage states, which in turn dictate the balance between fibrotic repair and tissue regeneration.

In this study, we address this gap by combining single cell and spatial transcriptomics with niche-aware computational modelling in the regenerating zebrafish heart. We map the composition and organisation of cardiac-immune microniches across homeostasis and regeneration, define how macrophage states distribute across these niches and identify niche-embedded signalling modules that couple macrophage identity to epicardial, vascular and fibroblast programmes. Mechanistically, we show that a fibroblast-driven il34 microniche programmes csf1ra^+^ macrophages into an egr1-dependent pro-regenerative state that restrains fibrotic programmes and supports vascularisation of the injury site. By identifying discrete ligand-receptor circuits and macrophage states that favour successful regeneration, our work provides entry points for re-engineering the mammalian cardiac niche away from scarring and towards regeneration.

## RESULTS

### Single cell profiling identifies discrete mpeg1.1+ immune populations in homeostatic and regenerating zebrafish hearts

To map the immune landscape of the adult zebrafish heart during regeneration, we isolated mpeg1.1^+^ cells from homeostatic, sham-operated and cryoinjured ventricles at 5 days post injury (dpi), a timepoint at which both pro-inflammatory and anti-inflammatory signatures coexist (8), and profiled these by single cell RNA sequencing (Figures 1A). The integrated dataset comprised 5,938 cells and resolved 21 transcriptionally distinct clusters, providing a coherent framework for condition-wise comparisons (Figures 1B,B^*′*^). We were able to capture the diversity expected within the mpeg1.1^+^ compartment in zebrafish (9), in which macrophages predominate and coexist with other innate and adaptive immune lineages (20).

**Figure 1.**
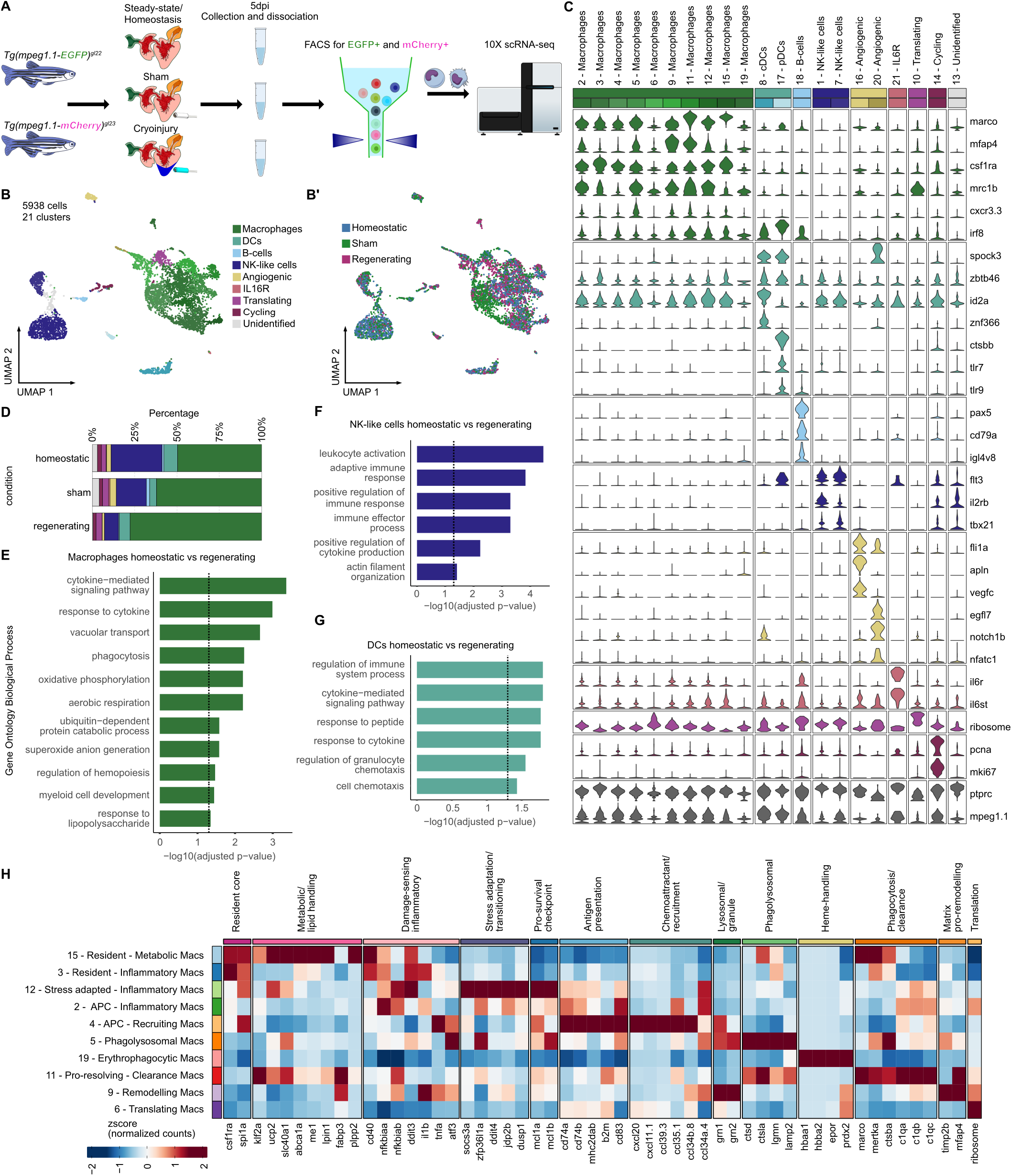
mpeg1.1^+^ immune cell populations in homeostatic and regenerating zebrafish hearts. A, Experimental design showing *Tg(mpeg1:EGFP)*^*gl22*^ and *Tg(mpeg1:mCherry)*^*gl23*^ reporters used for fluorescence-activated cell sorting (FACS) of mpeg1.1^+^ cells and single cell RNA sequencing. B,B*′*, UMAP embedding of 5,938 mpeg1.1^+^ cells coloured by transcriptional cluster (B) and by condition (B*′*), resolving macrophages, dendritic cells (DCs), B-cells, NK-like cells and minor populations. C, Violin plots of cell type-associated markers used to assign major immune identities and identify mpeg1.1^+^ DC, B-cell and NK-like subsets. D, Proportion of each mpeg1.1^+^ cell type across homeostatic, sham and cryoinjured hearts, showing a macrophage-centred immune environment in regenerating hearts. E-G, Gene Ontology enrichment analysis for differentially expressed genes in macrophages (E), NK-like cells (F) and DCs (G) in homeostatic versus regenerating hearts, highlighting activation of oxidative metabolism, phagocytosis and cytokine-mediated signalling programmes. H, Heat map of curated functional gene modules across macrophage clusters, defining resident-inflammatory, antigen-presenting, phagolysosomal, stress-adapted, pro-resolving, metabolic and erythrocyte-clearing states.

We assigned identities using cell type-associated markers consolidated from zebrafish immune single cell resources (8, 9, 20, 24, 25). As anticipated, macrophages were the most abundant cell types identified and expressed marco, mfap4, csf1ra and mrc1b (10, 26) (Figures 1C-D). Interestingly, we identified mpeg1.1^+^ dendritic cells in the heart and were able to resolve conventional dendritic cells (cDCs) expressing transcripts such as id2a, znf366, spock3 and plasmacytoid DCs (pDCs) expressing ctsbb, tlr7, tlr9 (20, 27, 28). mpeg1.1^+^ B-cells expressed pax5, cd79a and immunoglobulin transcripts such as igl4v8 (29); while mpeg1.1^+^ NK-like cells expressed core regulators of NK maturation such as tbx21 and il2rb, while lacking T cell genes such as il7r, cd4-1 and cd8a/b (24) (Figure 1C and Figure S1A).

We next asked how injury altered the balance of cell types. Quantifying the representation of each mpeg1.1^+^ population across conditions showed that macrophages, which were already the predominant cell type at homeostasis, expanded further after cryoinjury, whereas NK like cells declined from homeostatic through sham to regenerating conditions (Figure 1D). DCs and B-cells remained comparatively stable (Figure 1D). This redistribution establishes a mpeg1.1^+^ macrophage-centred immune environment in the regenerating heart, consistent with the regenerative requirement for zebrafish cardiac macrophages and their specialised subsets (9, 22) and in agreement with mammalian injury models showing that macrophages constitute the major immune component during repair (14, 30–32).

To interpret the major shifts in cell type composition, we examined differentially expressed genes and pathway enrichments within the principal compartments (Figures 1E-G, Figures S1B-D, Supplementary Information 1). Macrophages upregulated gene programmes linked to oxidative metabolism, phagocytosis and immune activation in cryoinjured hearts, consistent with debris clearance and post-injury remodelling (Figure 1E, Figures S1B, Supplementary Information 2). NK-like cells showed enrichment for adaptive immune and cytokine-mediated response and were reduced in numbers after injury (Figures 1D,F, Figures S1C, Supplementary Information 2). DCs increased chemotaxis and cytokine-responsive signatures, suggesting active sampling of the injured myocardium (Figure 1G, Figures S1D, Supplementary Information 2). These patterns mirror zebrafish single cell studies that distinguish pro-inflammatory and pro-remodelling myeloid states and reinforce the view that macrophage-centred programmes coordinate a permissive regenerative niche (9, 10, 20, 22).

Finally, in addition to the highly proliferative and translationally active groups of cells, we also noted two minor clusters with angiogenic and endothelial features and a small il6r-expressing cluster, that resembled immune subsets previously detected within zebrafish mpeg1.1^+^ compartments (20) (Figures 1B-C). Cluster 16 exhibited an endothelial-like signature likely reflecting immune-adjacent states rather than bona fide mpeg1.1^+^ immune cells. The haemogenic endocardial signature (notch1b, nfatc1, egfl7) was consistent with reports that the injured endocardium transiently acquires haemogenic/endothelial-to-haematopoietic transition features (33), and that, during development, the endocardium can generate myeloid cells (34–37). We therefore retained these clusters for context but excluded them from downstream analyses of immune cell crosstalk and function.

### Cardiac macrophage programme diversity underpins regenerative competence

To resolve the macrophage programmes that support regeneration, we focused on the mpeg1.1^+^ macrophage populations identified in Figures 1B-C, consisting of ten transcriptionally distinct clusters that were represented across conditions (Figures 1C,H, Figure S1E, Supplementary Information 3). Guided by a curated literature search and gene-module analysis, we assigned concise, function-oriented names that reflect dominant modules and likely roles in the injured heart (Figure 1H). These clusters collectively encode the macrophage-related functions associated with cardiac injury (2, 3). We therefore described these subpopulations in a way that reflects likely functional specialisation states for resident surveillance and damage sensing, antigen presentation and chemokine-driven recruitment, matrix remodelling and degradative phagocytosis, stress adaptation and resolution, metabolic/lipid handling and erythrocyte/heme clearance (Figure 1H, Figure S1E).

We first examined resident macrophage states. Cluster 15, “resident-metabolic”, co-expressed the resident core programme csf1ra, spi1a with a metabolic/lipid-handling module including klf2a, ucp2, slc40a1, abca1a, me1, lpin1, fabp3 and plpp2a, indicating resident macrophages tuned for lipid efflux and mitochondrial/redox metabolism. Cluster 3, “resident-inflammatory”, also carried the resident core genes but overlaid them with the damage-sensing inflammatory module expressing cd40, il1b, tnfa, nfkbiaa/b, ddit3 and atf3. This combination is consistent with bona fide tissue resident macrophages that act as early damage sensors and cytokine producers upon injury, rather than newly recruited monocytes. Together, these two resident populations delineate homeostatic versus activated resident programmes within the cardiac niche.

Among the more inflammatory states, Cluster 12, “stress adapted-inflammatory”, retained the damage-sensing inflammatory module but was further enriched for stress adaptation/transitioning genes such as socs3a, zfp36l1a, ddit4, jdp2b, dusp1 and the pro-survival checkpoint module mcl1a, mcl1b. This profile suggests macrophages that are adapting from acute cytokine production towards a more controlled, persistent phenotype. Cluster 2, “APC-inflammatory”, also encoded the damage-sensing inflammatory programme (cd40, il1b, tnfa, atf3, nfkbiaa/b, ddit3) but, in addition, up-regulated an antigen-presentation module including cd74a/b, mhc2dab, b2m, cd83. These features fit macrophages that bridge early innate cytokine production with MHC-II–mediated communication to lymphocytes, providing a natural link between inflammatory and antigen-presenting states.

Building on this, Cluster 4, “APC-recruiting”, shared the antigen-presentation module (cd74a/b, mhc2dab, b2m, cd83) but was distinguished by a prominent chemokine programme (cxcl20, cxcl11.1, ccl39.5, ccl35.1, ccl34a.4, ccl34b.8). This combination is consistent with macrophages acting as a chemoattractant antigen-presenting hub at the injury site, attracting additional leukocytes while presenting antigens.

We next identified phagocytic and cargo-handling states. Cluster 5, “phagolysosomal”, co-expressed the lysosomal/granule module grn1, grn2 and the phagolysosomal genes ctsd, ctsla, lgmn, lamp2, defining macrophages with high degradative capacity for damaged cardiomyocytes, matrix fragments and cellular debris. Cluster 19, “erythrophagocytic”, selectively expressed the heme-handling module (hbaa1, hbba2, epor, prdx2), a signature of red blood cells’ heme uptake and oxidative-stress buffering, consistent with erythrophagocytosis in haemorrhage- or vessel-adjacent regions, analogous to hbaa^+^ cardiac macrophages described in regenerative contexts (9).

Cluster 11 “pro-resolving-clearance” was dominated by phagolysosomal genes (ctsd, ctsla, lgmn, lamp2) together with a phagocytosis/clearance programme (marco, mertka, ctsba, c1qa, c1qb, c1qc). The combination of scavenger receptors, complement components and degradative machinery is typical of efferocytic, pro-resolving macrophages that clear apoptotic cells and debris while supporting dampening of inflammation (9, 20, 38). Cluster 9 “remodelling” was enriched for the matrix pro-remodelling module timp2b, mfap4, alongside lysosomal/granule genes grn1, grn2, indicating macrophages focused on ECM turnover and processing of engulfed material at the injury site. These two clusters therefore represent complementary late-phase programmes of clearance and structural remodelling that are already present at 5dpi.

Finally, Cluster 6, “translating”, was marked by increased expression of ribosomal genes and displayed mixed functional signatures, consistent with a transcriptionally heterogeneous but translationally engaged macrophage state that may underpin dynamic switching between the programmes described above.

Together, these co-existing programmes reveal a diverse macrophage landscape in which resident and recruited cells function distinctly across damage sensing, leukocyte recruitment, regulation and resolution of inflammation, metabolic support and erythrocyte clearance, underpinning the regenerative competence of the injured zebrafish heart.

### Spatial mapping and ligand-receptor-target inference identify candidate niche-macrophage signals

To connect macrophage subpopulations to their broader tissue context and identify candidate regulatory signals that might be shaping macrophage diversity and function, we first performed spatial transcriptomics using the 10x Visium platform on homeostatic and cryoinjured ventricles (Figures 2A-C). We deconvoluted Visium spots with cell2location (39), using our mpeg1.1^+^ single cell atlas complemented with whole-heart reference datasets (40), to infer the contribution of major structural and immune cell types to each Visium spot. As predicted, a large number of smooth muscle cells localised to the outflow tract, while cardiomyocytes were mapped to the ventricular wall, with fibroblasts and epicardial cells residing in the outer layer of the heart (Figures 2D,D^*′*^). In contrast, macrophages were restricted to sparse epicardial and perivascular regions in the homeostatic heart, becoming more concentrated around the injury region and the epicardial layer in the regenerating heart (Figure 2D^*′*^). Although this spot-based mapping confirmed gross anatomical patterns, the effective resolution of Visium, each spot pooling transcripts from several neighbouring cells, was insufficient to resolve individual macrophage states or to model cardiac niche-macrophage crosstalk with the required granularity to determine how immune and stromal cells are spatially organised and connected during cardiac regeneration.

**Figure 2.**
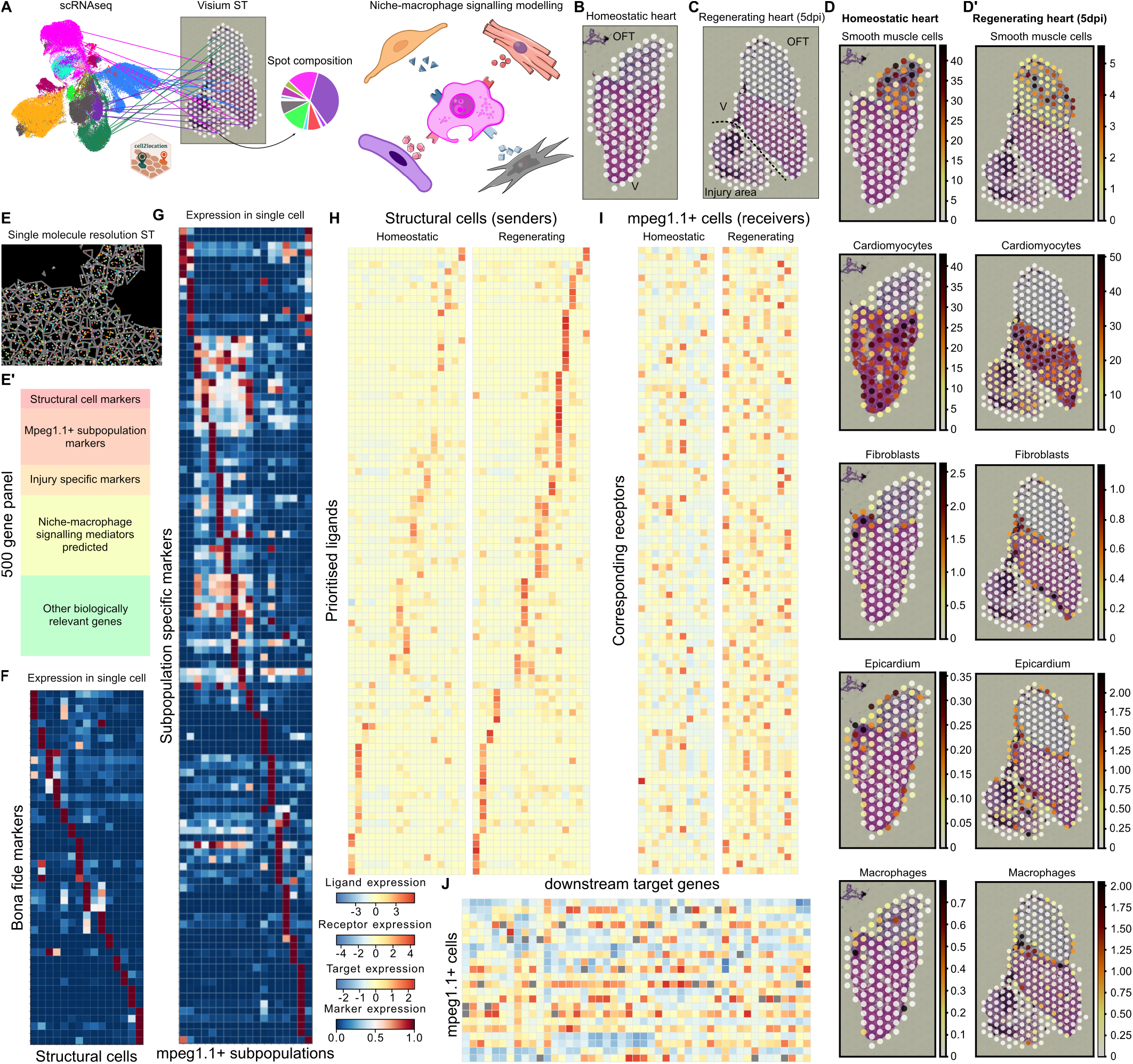
Visium-NicheNet pipeline and MERFISH panel design to prioritise candidate niche-macrophage signals. A, Schematic of the integrative analysis pipeline combining single cell RNA-seq, Visium spatial transcriptomics and ligand-receptor-target inference to identify niche-macrophage signalling programmes. B,C, Visium sections from homeostatic (B) and 5 dpi regenerating (C) hearts, with each white spot representing a 55µm capture area. D,D*′*, Spatial maps of key structural cell types identified (smooth muscle cells, cardiomyocytes, fibroblasts, epicardium and macrophages) in homeostatic (D) and regenerating (D*′*) hearts according to inferred cell type composition after cell2location deconvolution. E, Representative MERFISH image illustrating single molecule transcript detection and cell segmentation. E*′*, Heat map summarising the biological categories included in designing the 500 gene MERFISH panel, grouped into structural cell markers, mpeg1.1^+^ subpopulation genes, injury-induced signatures, candidate niche-macrophage signalling mediators and other regeneration-related genes. F,G, Marker selection matrices used to distinguish structural cells (F) and mpeg1.1^+^ immune subpopulations (G) in MERFISH data. Detailed gene panel design included in Table S1. H-J, NicheNet-based prioritisation of ligand-receptor circuits, showing ligands upregulated after injury in structural “sender” populations (H), corresponding receptor activity in mpeg1.1^+^ “receiver” subsets (I) and predicted downstream target genes in mpeg1.1^+^ macrophages (J) that together define putative functional communication programmes. Detailed NicheNet output included in Supplementary Information 4.

We therefore used these data as a scaffold to design a higher-resolution MERFISH (41) experiment using the imaging-based spatial transcriptomics Merscope platform that allows for single molecule-resolution capture of transcripts *in situ* (Figure 2E). We assembled a 500-gene panel that combined five classes of genes: bona fide markers distinguishing structural cell types that make up the cardiac niche, markers that uniquely identify the mpeg1.1^+^ subpopulations from our single cell transcriptomics data (Figure 1), injury-induced genes, predicted niche-macrophage signalling mediators, as well as key regeneration-related signatures (Figures 2E-G). Description of the 500 gene panel design included in Table S1. To enrich the panel for functional communication circuits, we adapted the NicheNet pipeline (42) for the zebrafish genome, treating structural cells as putative “senders” and mpeg1.1^+^ subpopulations as “receivers”. This analysis allowed us to prioritise a restricted set of ligands upregulated after injury in structural compartments (Figure 2H), infer corresponding receptor activity on mpeg1.1^+^ cells in regenerating versus homeostatic hearts (Figure 2I) and predict coherent modules of downstream target genes within macrophages as putative effectors mediating their identity and function (Figure 2J). We further curated this list by literature-based filtering, retaining ligands, receptors and target genes with clear links to immune regulation, tissue remodelling and regeneration. Detailed NicheNet output of prioritised ligands, corresponding receptors and downstream targets was included in Supplementary Information 4.

Together, the Visium-NicheNet pipeline provided both a coarse spatial framework and a mechanistically informed gene set, nominating concrete hypotheses for niche-macrophage communication to be tested in subsequent MERFISH experiments.

### Spatial cardio-immune microniches with distinct regulatory programmes emerge during early regeneration

To resolve fine-grained cellular neighbourhoods and decode the molecular programmes that mediate niche-macrophage signalling during heart regeneration, we next performed MERFISH on homeostatic (Figure 3A) and 5 dpi regenerating (Figure 3B) zebrafish hearts. Cell type label transfer (43) using our single cell reference revealed a marked expansion of macrophages and fibroblasts within the injured region, accompanied by depletion of cardiomyocytes (Figures 3A^′^,A^′′^,B^′^,B^′′^,C). These changes mirrored those observed in dissociated single cell suspension tissues (40, 44, 45) and confirmed early immune and mesenchymal remodelling at the lesion site.

**Figure 3.**
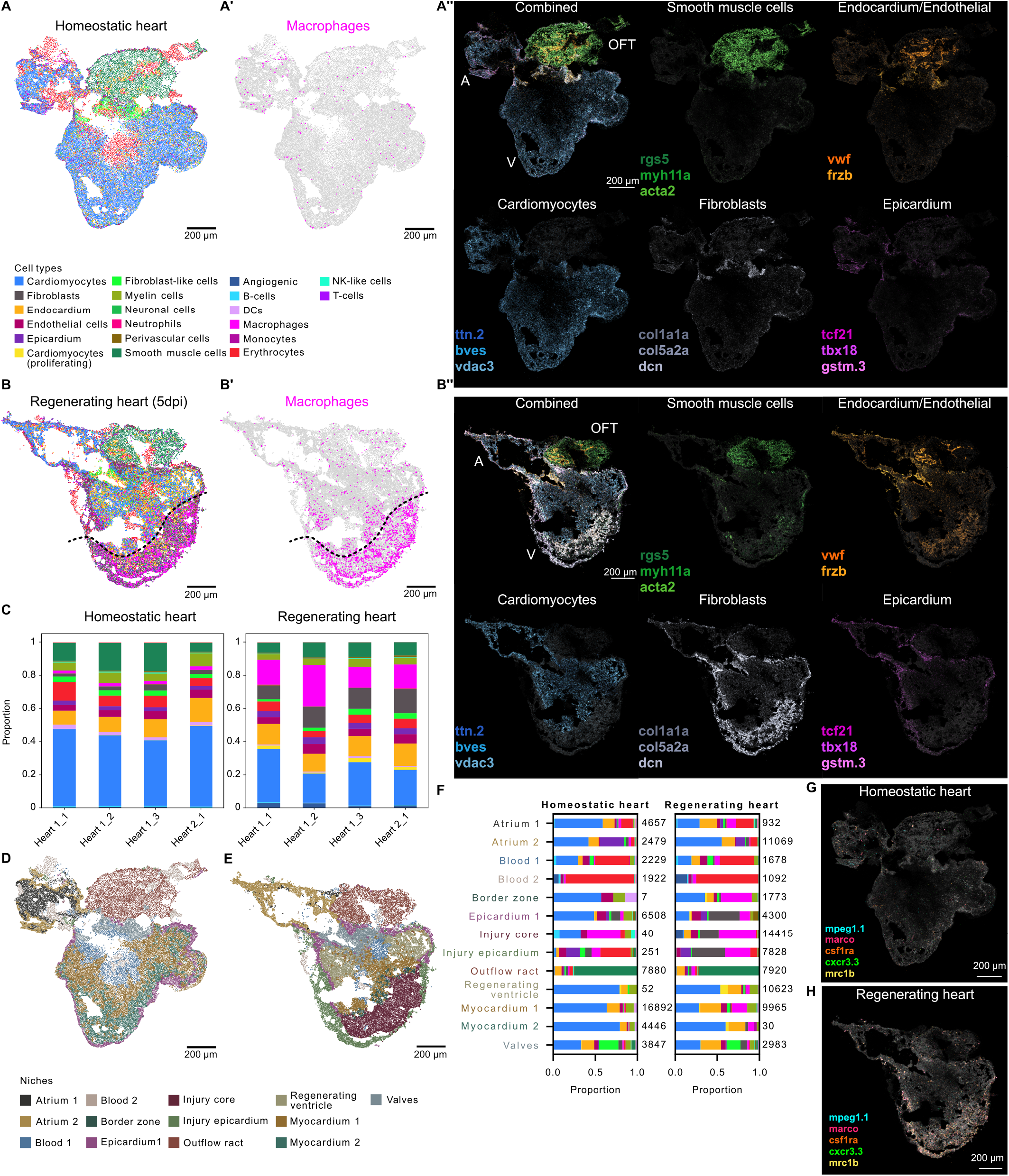
Spatial mapping of cell types and niches in homeostatic and regenerating hearts. A,A*′*, MERFISH maps of homeostatic hearts with cells coloured by cell type (A) and with macrophages highlighted (A*′*). A*′′*, MERFISH spatial expression of representative cell type markers (smooth muscle cells, endocardium/endothelium, cardiomyocytes, fibroblasts and epicardium) in homeostatic hearts. B,B*′*, Maps of 5 dpi regenerating hearts coloured by cell type (B, cell type key as in A), highlighting macrophage distribution (B*′*). B*′′*, MERFISH spatial expression of cells type markers in regenerating hearts, illustrating depletion of cardiomyocytes and upregulation of fibroblast and macrophage signatures in the injury region. C, Proportion of structural and immune cell types per section (n=4 per condition) across biological replicates (n=2 per condition) in homeostatic and regenerating hearts, cell type key as in A. D,E, NicheCompass-derived multicellular niches in homeostatic (D) and regenerating (E) hearts, capturing anatomical domains including atrium, valves, myocardium, regenerating ventricle, border zone, injury core, injury epicardium, blood and outflow tract. F, Bar plots showing stereotyped cell type numbers and composition of each niche in homeostatic and regenerating conditions, cell type key as in A. Spatial distribution of macrophages in the homeostatic (G) and regenerating (H) hearts, with spatial expression of the macrophage markers mpeg1.1, marco, csf1ra, mrc1b and cxcr3.3 distributed along the injured heart, including the injury core and injury epicardium niches.

To spatially resolve this response, we applied NicheCompass (46), a communication-aware framework that segments tissues into discrete multicellular domains using ligand-receptor and signalling pathway embeddings. This analysis delineated reproducible domains across the homeostatic (Figure 3D) and regenerating heart (Figure 3E). Each niche domain exhibited a stereotyped cell composition and spatial localisation, consistent with anatomical patterning and emergent multicellular niches, encompassing the epicardium, injury core, border zone, myocardium, atrium, valves and the outflow tract (Figure 3F). We identified myeloid cells (Figure 3G), which formed a noticeable border at the lesion interface, with macrophages expressing mpeg1.1, marco, csf1ra, cxcr3.3 and mrc1b enriched along the injury area (Figure 3H). This accumulation outlined the regenerating zone, including the regenerating ventricle, injury core, injury epicardium and border zone niches (Figure 3D), reflecting principles of local immune enrichment described in zebrafish and mammalian infarct models (3, 14, 16).

We hypothesised that these injury-based macrophages were not uniformly activated but instead shaped by microanatomical cues. Therefore, to investigate if discrete microenvironments organise macrophage-structural cell interactions, we trained a dedicated NicheCompass model to infer interaction programmes in the regenerating injury area only (Figure 4A), where most of the macrophage subpopulations resided (Figure 3H). Here, we identified multiple discrete microenvironments, or finer-grained microniches (Figure 4A^*′*^), rather than a uniform signalling landscape, each characterised by a distinct pattern of potential cell-cell communication gene programmes (GPs) (Figure 4A^*′*^, Figure S2A). Specifically, each microniche exhibited a unique combination of cell populations acting as potential dominant signal senders (Figure 4B), which led us to next investigate if distinct macrophage subpopulations (as defined in Figure 1) could be serving as the primary signal receivers. We first found that these GP signalling circuits were highly spatially confined and directionally coherent, meaning that specific cell types in defined microniches sent signals to neighbouring target cells in an organised way (Figures 4C-F, Figure S2A).

**Figure 4.**
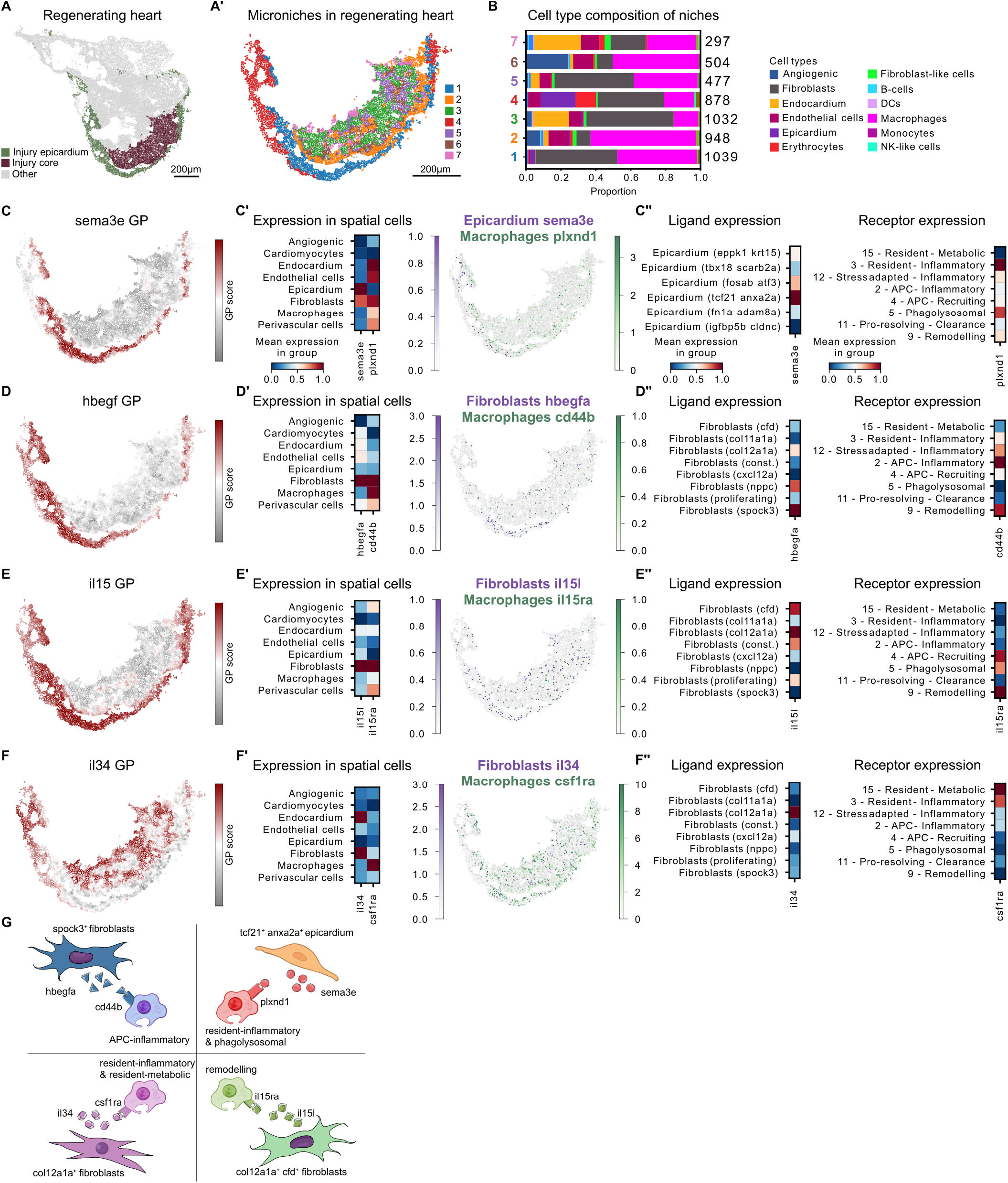
Distinct microniche gene programmes instruct macrophage states in the injury zone. A, NicheCompass map of the regenerating heart at 5 dpi with the injury core and injury epicardium highlighted. A*′*, NicheCompass segmentation of the injury territory into microniches (1-7), each representing a discrete microenvironment. B, Cell type composition of each microniche, revealing distinct combinations of epicardium, fibroblasts, endothelial cells, cardiomyocytes and macrophage populations. C–F, Representative ligand-receptor gene programmes enriched in selected microniches. C,C*′*,C*′′*, Sema3e programme: spatial distribution of the sema3e gene programme score (C), expression of sema3e and plxnd1 across major structural cell types and macrophages (C*′*) and assignment of sema3e and plxnd1 expression to epicardial and macrophage subclusters using single cell data (C*′′*). D,D*′*,D*′′*, Hbegf programme: spatial distribution of the Hbegf gene programme score (D), expression of hbegfa and cd44b across structural and immune populations (D*′*) and mapping of sender spock3^+^ fibroblasts and receiver APC-inflammatory macrophage subclusters (D*′′*). E,E*′*,E*′′*, Il15 programme: spatial pattern of the Il15 gene programme score (E), expression of il15l and il15ra across cell types (E*′*) and localisation of il15l^+^ col12a1a^+^/cfd^+^ fibroblasts and il15ra^+^ macrophage subclusters (E*′′*). F,F*′*,F*′′*, Il34-csf1ra programme: spatial distribution of the Il34 gene programme score (F), localisation of il34 expression to col12a1a^+^ fibroblasts and csf1ra expression to macrophages (F*′*) and assignment of il34 senders and csf1ra^+^ resident macrophage receivers at single cell resolution (F*′′*). G, Schematic summarising the main macrophage-centric signalling circuits identified, illustrating how distinct microniches couple structural cell programmes to specialised macrophage states.

We then focused on representative microenvironment regions and their dominant ligand-receptor circuits. Microniches 1 and 4 comprised the epicardial layer and immediately subjacent sub-epicardial fibroblasts (Figures 4A^′^,C), hosting signalling programmes that are spatially restricted to the outer layer of the lesion. We first focused on the sema3e GP by analysing our spatial transcriptomics data and found epicardial cells as the dominant senders expressing this guidance cue molecule, with macrophages within the same neighbourhood being enriched for plxnd1 receptor transcripts (Figure 4C^′^). To assign sender and receiver subpopulations with higher granularity, we correlated our single cell suspension data with our Merscope spatial maps to assign expression of these GP mediators to specific cellular subsets. By doing so, we identified a subpopulation of epicardial cells characterised by increased expression of tcf21 and anxa2a, forming the sender population with marked expression of sema3e. At the receiver side, cluster 3 “resident-inflammatory” macrophages, which combine the resident core and damage-sensing inflammatory modules, as well as cluster 5 “phagolysosomal” macrophages, enriched for lysosomal/granule and phagolysosomal programmes, exhibited higher levels of plxnd1 expression (Figure 4C^*′′*^). Sema3e has been implicated in regulating macrophage distribution in hypoxic tissue zones, thus, the epicardial sema3e microniche signal could help organise a regenerative microenvironment, possibly attracting or positioning resident-derived inflammatory and phagolysosomal macrophages in the sub-epicardial layer to facilitate regeneration of the damaged tissue (47). Within the same microniches, we then focused on the hbegf GP (Figure 4D), given their link to cardiac regeneration (48). We found the ligand heparin-binding EGF-like growth factor, hbegfa, to be highly expressed in spock3^+^ fibroblasts, with its receptor cd44b expressed in the adjacent macrophage subpopulation 2 “APC-inflammatory” macrophages (Figures 4D^*′*^,D^*′′*^), which couple the damage-sensing inflammatory and antigen-presentation modules.

We next studied the core injury territory by focusing on the remaining microniches and the related fibroblast-derived programmes. We found the il15 GP (Figure 4E) to be dominantly represented by fibroblasts as sender cells expressing il15l (Figure 4E^*′*^). At single cell resolution, expression of this ligand mapped to col12a1a^+^ and cfd^+^ fibroblasts (Figure 4E^*′′*^). Neighbouring receivers included macrophage subclusters expressing the Il-15 receptor il15ra (Figure 4E^*′′*^) namely cluster 9 “remodelling” macrophages, which express the matrix pro-remodelling and lysosomal/granule programmes. Finally, the il34 GP (Figure 4F) identified a discrete col12a1a^+^ subpopulation of fibroblasts expressing the ligand il34 as the main sender cells (Figures 4F^*′*^,F^*′′*^) and csf1ra^+^ resident-derived macrophage states, including clusters 3 “resident-inflammatory” and 15 “resident-metabolic” as receiving cells within the same microniche. The downstream target module is centred on activation of the transcription factor egr1, expressed in most of the macrophage subpopulations identified, with enriched expression detected in cluster 11 “pro-resolving-clearance” macrophages, which carry the phagolysosomal and phagocytosis/clearance programmes (Figures 4F^*′*^,F^*′′*^, Figures S2B,B^*′*^). This il34-csf1ra-egr1 axis connects fibroblast senders to resident and pro-resolving macrophage receivers.

These programmes underscore the role of the local environment in shaping macrophage transcriptional identity, with structural-immune signalling circuits driving microniche-specific macrophage phenotypes (Figure 4G).

### Fibroblast-macrophage microniche triggers egr1 in macrophages through il34-csf1ra signalling

Building on our spatial transcriptomic analysis of GPs in regenerating hearts (Figure 4), we next functionally tested the il34-csf1ra-egr1 signalling module (Figure 5). To specifically disrupt this macrophage activation circuit, we analysed the csf1ra^j4e1/j4e1^ zebrafish, which lack functional Csf1ra receptors and represent a well-established model of altered macrophage-dependent cardiac repair (10, 21) (Figures 5A,A^*′*^, Figures S3A,A^*′*^). At the tissue-wide level, csf1ra^j4e1/j4e1^ hearts showed preserved infiltration of macrophages within the injury zone (Figure 5A^*′*^) but displayed markedly altered cellular composition when compared with their wild type counterparts, particularly within the immune compartment (Figure 5B). Spatial label transfer revealed an overall reduction in macrophages, endothelial and epicardial cells and an increase in fibroblasts in the injury territory compared to wild type (Figure 5B). While global niche structure was retained relative to wild type (Figures 5C,C^*′*^, Figures 3D,E, Figure S3B), finer-grained spatial mapping showed that the regenerative architecture was subtly disrupted in the mutant heart (Figure 5D, Figure 4A^*′*^). We observed a selective expansion of microniches 5 and 6 (Figure 5D, Figure S3C), and reduction of microniches 1, 3 and 4 (Figure 5D, Figure S3C).

**Figure 5.**
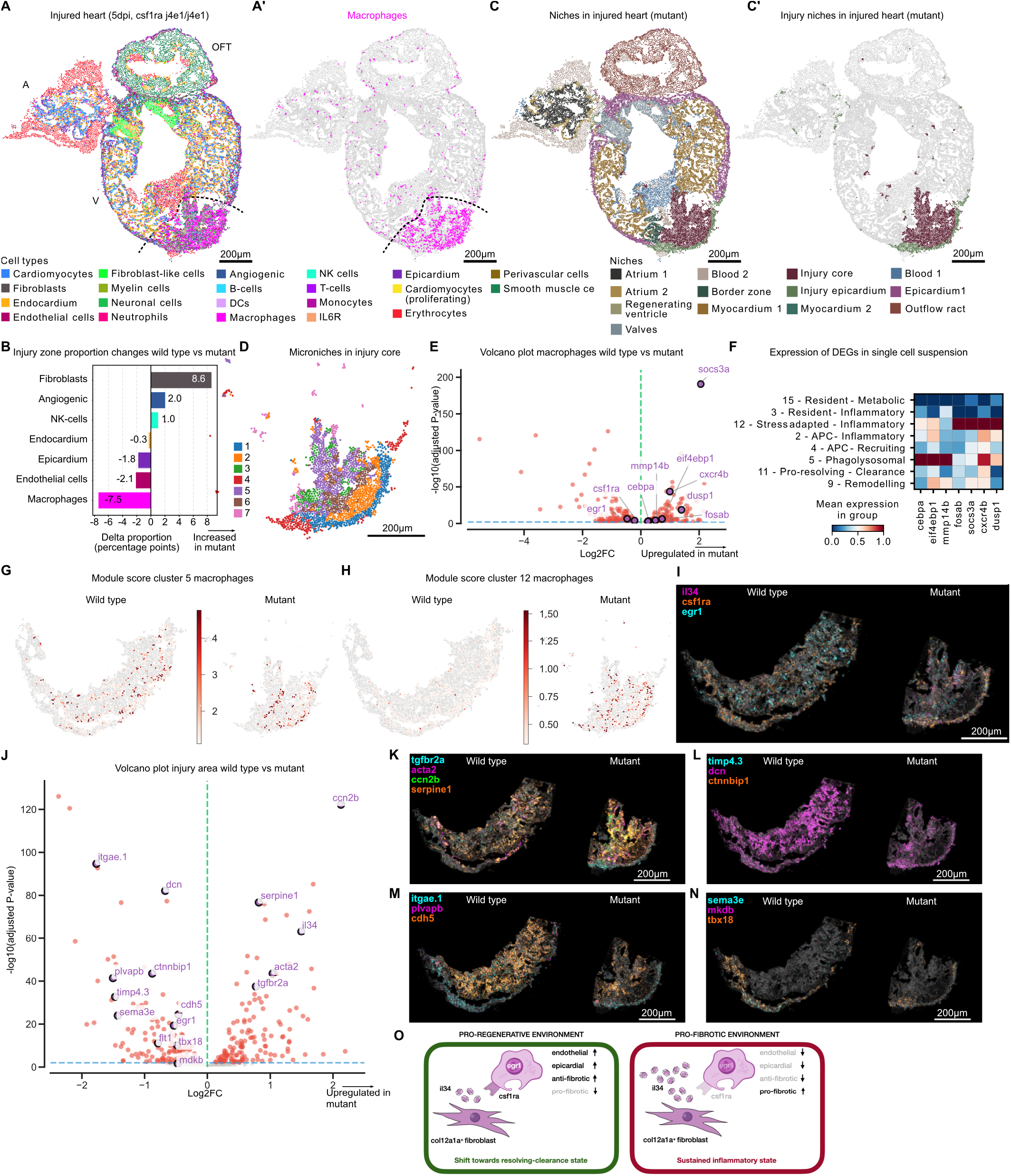
Disruption of the il34-csf1ra-egr1 axis remodels fibroblast-macrophage microniches. A,A*′*, MERFISH maps of the injured heart of csf1ra^j4e1/j4e1^ mutants at 5 dpi, coloured by cell type (A) and highlighting macrophages (A*′*). B, Comparison of cell type proportions in the injury territory between wild type and csf1ra mutants. C,C*′*, NicheCompass-defined niches in the mutant heart (C) and injury-restricted niches (C*′*). D, Microniches within the mutant injury region. E, Volcano plot of differentially expressed genes in macrophages in wild type compared to mutant injury territories. F, Heatmap of genes upregulated in mutant macrophages and ploted across macrophage clusters, showing preferential enrichment of cluster 5 “phagolysosomal” and cluster 12 “stress-adapted inflammatory” signatures in mutants. Spatial module scores for clusters 5 (G) and 12 (H) projected onto wild type and mutant hearts, illustrating selective upregulation of programmes and increase of stress-adapted inflammatory states, particularly cluster 12, in mutant heart. I, Spatial expression of il34, csf1ra and egr1 in wild type and mutant hearts within the injury zone. J, Volcano plot of differentially expressed genes within the injury territory of wild type versus csf1ra mutants. K-N, MERFISH spatial expression maps of fibrosis-related genes tgfbr2a, acta2, ccn2b, serpine1 (K), ECM remodelling genes dcn4, timp4.3, ctnnbip1 (L), endothelial factors itgae.1, cdh5, plvapb (M) and epicardial factors sema3e, mdkb, tbx18 (N) in the wild type and mutant injury area. O, Model summarising how disruption of il34-csf1ra signalling prevents egr1 induction in resident and pro-resolving macrophages, shifting macrophages towards stress-adapted inflammatory states, which impacts endothelial and epicardial compartments and biases repair towards fibrosis.

In csf1ra mutant, macrophages upregulated genes such as cebpa, eif4ebp1, mmp14b, fosab, socs3a, cxcr4b, and dusp1 (Figure 5E). We noted that many of these upregulated signatures mapped onto transcriptional states corresponding to cluster 5 “phagolysosomal” and cluster 12 “stress adapted-inflammatory” macrophages of our single cell transcriptomics data (Figure 5F). Cluster 5 was characterised by co-enrichment of the lysosomal/granule and phagolysosomal programmes, whereas cluster 12 combined the damage-sensing inflammatory module with stress adaptation/transitioning and pro-survival checkpoint genes (Figure 1H). In accordance, we detected an expansion of these signatures when spatially mapped onto the mutant heart (Figures 5G,H), with expansion of subpopulation 12 being much more marked. This suggests that loss of csf1ra signalling selectively amplifies macrophage states associated with phagolysosomal activation and stress-adapted inflammatory programmes.

We next examined expression of the il34-csf1ra-egr1 signalling components in wild type and mutant injured hearts (Figure 5I, Figures S3D,E). We detected expression of the ligand il34 in fibroblasts marked by col1a1a and egr1 transcripts colocalising with csf1ra-expressing macrophages at the site of injury of wild type hearts. In contrast, expression of the downstream transcription factor egr1 was markedly reduced in csf1ra^j4e1/j4e1^ mutants in the injury zone (Figure 5I, Figures S3D,E). Interestingly, fibroblast-niche ligand expression showed compensatory il34 upregulation in mutant hearts (Figure 5I, Figures S3D,E), suggesting a feedback attempt to stimulate macrophages. However, this ligand compensation was insufficient to rescue egr1 induction, indicating a requirement for functional receptor activation. Notably, although csf1ra transcripts were still detected in mutants, consistent with the csf1ra^j4e1/j4e1^ being hypomorphic rather than a full null (21), this did not translate into functional signalling, as il34 failed to trigger egr1 expression (Figure 5I, Figures S3D,E). Differential expression analysis confirmed egr1 downregulation in mutant versus wild type injury regions (Figure 5J). Importantly, egr1 suppression was spatially restricted to the injury territory, as confirmed by analysis of non-injured zones (Figures S3F,G,H), demonstrating that this il34-csf1ra-egr1 axis disruption is not due to systemic transcriptional repression but rather reflects a microniche-related defect. This finding directly confirms the NicheCompass-inferred circuit from our spatial analysis (Figure 4F), demonstrating that fibroblast-derived il34 engages csf1ra^+^ resident and pro-resolving-clearance macrophage subsets to trigger an activation state marked by egr1 and associated genes. Such microniche-induced transcriptional switch towards egr1-high pro-resolving-clearance and resident programmes seems essential for coordinating regenerative fibroblast-macrophage interactions at the injury site.

In accordance, across the injury area of mutant hearts we observed upregulated signalling linked to fibrosis, shown by increased expression of pro-fibrotic signatures such as ccn2b, serpine1, acta2, tgfbr2a (Figures 5J,K) and reduced expression of remodelling genes such as dcn, timp4.3 and ctnnbip1 (Figures 5J,L). Strikingly, we noted that vascular integrity factors such as cdh5, plvapb and itgae.1 were downregulated, suggesting a destabilised endothelial niche in csf1ra mutants (Figures 5J,M). Furthermore, we also noted epicardial factors such as tbx18, sema3e and mdkb to be downregulated in the mutant injury area, indicating a disruption of the epicardial signalling compared to wild type (Figures 5J,N). Such endothelial and epicardial signature-specific disruption was in accordance with our NicheCompass analysis (Figure 5B).

In summary, fibroblast-derived Il34 normally signals via Csf1ra on injury-site macrophages, particularly resident-derived and pro-resolving-clearance states, to induce an egr1-high, pro-regenerative environment that restrains fibrosis and supports vascular integrity and epicardial signalling. In csf1ra mutants, this axis is uncoupled, il34 is upregulated but cannot signal effectively, macrophages shift towards phagolysosomal and prolonged stress adapted-inflammatory states, fibroblasts domains expand, and endothelial and epicardial identities are compromised, collectively biasing the early injury response towards a pro-fibrotic repair programme (Figure 5O).

To make our MERFISH data accessible to the community, we developed an interactive dashboard that provides access to our processed Merscope data. The application is hosted at: https://simoeslab-regenerating-heart-dashboard.hf.space/

## DISCUSSION

Our study reveals that cardiac regeneration in the adult zebrafish is coordinated by a macrophage compartment comprising diverse sub-populations spatially confined within distinct cardio-immune microniches. By combining single cell transcriptomics, computational ligand-receptor inference and imaging-based spatial transcriptomics, we define a macrophage-centred immune landscape and identify a fibroblast il34-csf1ra-egr1 pathway as a key pro-regenerative axis. Together, these data shift emphasis from a largely population-averaged view of “regenerative-prone macrophages” to a niche-centred framework in which cardiac structural cells instruct distinct macrophage states with specialised functions.

A central aspect of our work is the mechanistic dissection of how these spatial programmes shape a spectrum of resident, inflammatory, antigen-presenting, pro-resolving and remodelling macrophage states. These states are evident in the single cell suspension data but acquire clearer functional relevance when mapped into their cardiac microniches. This in turn provides a mechanistic explanation as to how macrophages can simultaneously execute debris clearance, antigen presentation, matrix remodelling and resolution within a relatively small, injured territory, as opposed to adopting a single averaged activation state. It also suggests that the regenerating heart is instructed by a small number of recurring structural-immune building blocks, in line with the concept of communication-defined niches emerging from NicheCompass and related frameworks (46).

Our findings build on previous work implicating Csf1ra in macrophage-dependent scar formation and regeneration in the zebrafish heart (10, 21). However, herein we reveal that Csf1ra does not simply control macrophage numbers or a TNFα^+^ subset, but instead acts as a receiver of col12a1a^+^ fibroblast-derived il34 to establish an egr1^+^ pro-resolving-clearance and resident macrophage programme within a specific microniche. Loss of csf1ra skews the early cardiac immune landscape towards prolonged stress-inflammatory states and is associated with early pro-fibrotic programming and destabilisation of vascular and epicardial niches. By combining spatial deconvolution with state-resolved mapping, we now show that loss of Csf1ra selectively erodes the fibroblast-macrophage microniche and, as a consequence, resident-inflammatory macrophages fail to occupy their microenvironment leading to egr1 downregulation. In this scenario, il34-rich fibroblast microniches expand at the expense of macrophage-, endothelial- and epicardial-rich domains. In parallel, phagolysosomal and stress adapted-inflammatory macrophages become over-represented and upregulate stress and inflammatory programmes, while genes associated with pro-resolving-clearance are reduced. These observations reconcile previously divergent phenotypes by showing that Csf1ra is required not only for macrophage abundance, but also for sustaining a pro-regenerative spatial configuration of macrophage states.

Our data further place the il34-csf1ra-egr1 axis as a key molecular switch within this configuration. In csf1ra mutants, il34 expression is paradoxically increased, yet fails to induce egr1, confirming that intact receptor-driven macrophage signalling is required for this niche-induced programme. This fits with broader evidence that stromal niches use CSF1 and IL-34 to sustain tissue resident macrophages across organs, creating “two-cell circuits” in which fibroblasts, endothelial cells or epithelial cells provide trophic CSF1R ligands to nearby macrophages (49). Work in human macrophages has shown that EGR1 acts as a gatekeeper of inflammatory enhancers, repressing pro-inflammatory gene programmes and blunting excessive activation while helping to establish macrophage identity through CSF1R-linked regulatory elements (50). Our findings extend these concepts *in vivo* by showing that, in the regenerating heart, a fibroblast-derived il34 signal uses csf1ra to activate an egr1^+^ pro-resolving-clearance macrophage state within a defined microniche. The absence of egr1^+^ macrophages in csf1ra^j4e1/j4e1^ injury microniches likely underlies the disordered inflammation and altered repair dynamics in these mutants (10), highlighting that Egr1 functions as a macrophage-intrinsic switch that restrains inflammatory activity while permitting regenerative functions. This is consistent with recent work in the mouse showing that EGR1 coordinates a regenerative senescence-response in cardiac fibroblasts and promotes angiogenesis and cardiomyocyte proliferation during repair (51).

The downstream consequences of losing this axis highlight how subtle defects in niche communication can tilt regeneration towards fibrotic repair. In csf1ra mutants at 5 dpi, pro-fibrotic markers (ccn2b, acta2) are upregulated while remodelling genes (dcn, timp4.3) are reduced, consistent with persistent myofibroblast activation and impaired matrix turnover. These early programme-level defects provide a mechanistic explanation for the residual scarring described in csf1ra-deficient hearts at 21 dpi (10) and suggest that the Il34-Csf1ra-Egr1 axis is required to drive fibroblasts and macrophages from a fibrotic into a resolving state. Endothelial and epicardial signatures are also downregulated, with reduced expression of vascular adhesion molecules and epicardial factors, indicating destabilisation of these structural niches. These changes resemble vascular-perivascular remodelling in mammalian heart failure and align with the idea that Vwf^+^ niches are critical for effective repair (16, 31, 52). In contrast, the zebrafish heart can normally re-establish vessels and resolve scar tissue after injury (4, 8, 53, 54) and our data suggest that maintaining il34-csf1ra-egr1-licensed macrophages within this microniche is one mechanism that keeps the system on a regenerative trajectory.

More broadly, our work aligns the zebrafish cardiac niche with emerging principles of spatially organised immune regulation across organs. Spatial multi-omics studies in lymphoid organs and barrier tissues have revealed that discrete cytokine microniches, for example, IL-2/IL-10 niches in lymph nodes and the lung stabilise local T cell fates and constrain effector functions to specific microenvironments (17, 18). Analogously, we find that epicardial and fibroblast-rich microniches use distinct ligand-receptor circuits, with Sema3e and Hbegfa signals from the epicardial/sub-epicardial layer engaging resident-inflammatory, phagolysosomal and APC-inflammatory macrophages, while Il15l and Il34 signals from injury-core fibroblasts are priming remodelling, resident-inflammatory, resident-metabolic and pro-resolving-clearance macrophages. The fact that NicheCompass, initially developed for embryonic and tumour tissues, cleanly resolves these cardiac microniches underscores the generalisability of communication-centric niche models (46).

Our findings also speak to the long-standing debate over whether cardiac macrophages promote regeneration or fibrosis. In mammals, macrophages can support angiogenesis and electrical conduction but also drive adverse remodelling, depending on context (31, 55).In zebrafish, perturbing macrophages or the vascular niche impairs regeneration (9, 22, 25, 56, 57). By stratifying macrophages into niche-encoded states, we show that pro- and anti-regenerative activities coexist within the same tissue and that the balance between remodelling, stress-adapted inflammatory, pro-resolving and metabolic states is spatially regulated. This explains why broad macrophage manipulation in mammals has produced mixed outcomes and argues for strategies that target sub-populations residing in specific niches rather than the entire macrophage compartment.

In conclusion, we establish that cardiac regeneration in zebrafish is orchestrated by a small set of spatially confined structural-immune microniches and that a fibroblast-driven Il34-Csf1ra-Egr1 axis is necessary to populate these microenvironments with pro-regenerative macrophages, while restraining persistent stress adapted-inflammatory programmes that lead to fibrotic repair and vascular destabilisation. This work bridges the gap between descriptive immune atlases and mechanistic niche biology, providing a template for using spatial multi-omics to decode communication logic in other regenerative and fibrotic settings. By identifying definitive ligand-receptor modules and macrophage states that correlate with regenerative outcome, we also highlight putative targets for re-engineering the mammalian cardiac niche towards regeneration rather than permanent scarring.

## METHODS

### Experimental Model and Subject Details

This study was carried out in accordance with procedures authorised by the UK Home Office in accordance with UK law (Animals Scientific Procedures Act 1986) and approved by the Research Ethics Committee of the University of Oxford. For this study, both females and males of wild type, transgenic and mutant zebrafish strains were used. Animals used for breeding were between 4 and 24 months old, kept at a 14 hours light, 10 hours dark cycle and fed four times a day.

### Zebrafish Lines

Published transgenic and mutant lines used in this study were *Tg(mpeg1:EGFP)*^*gl22*^, *Tg(mpeg1:mCherry)*^*gl23*^ (58) and the csf1ra^j4e1/j4e1^ panther mutant (10, 21).

### Zebrafish cardiac injury model

Cardiac injuries on adult zebrafish were carried out as previously described (53, 59). Briefly, zebrafish aged 6 to 18 months old were anesthetised in 1 g/L MS222 (E10521, Sigma-Aldrich) and placed ventral side facing up in a wet sponge with a slit. To first expose the heart, an incision was made through the pericardium using surgical scissors. To induce ventricular cryoinjury, the exposed ventricle was dried and a liquid nitrogen-cooled cryoprobe was applied to the surface of the ventricle until the probe was completely thawed to damage approximately 20% of the tissue. In sham control animals, an incision was made through the pericardium, and the exposed ventricle was gently touched with a probe at room temperature without causing injury. After surgery, the fish were returned to systems water and allowed to fully recover before being transferred back to the main aquarium system until the desired timepoint.

### Heart tissue dissociation and isolation of mpeg+ cells by FACS

19 sham *Tg(mpeg1:GFP)*^*gl22*^ and 22 cryoinjured hearts *Tg(mpeg1:mCherry)*^*gl23*^ were collected at 5 days post-surgery, together with 16 unopened *Tg(mpeg1:GFP)*^*gl22*^ hearts. The collected hearts were placed in ice-cold Ringer’s solution and transferred to 1X HBSS (14185052, Life Technologies). Hearts were dissociated into single cell suspensions at 32°C by incubating with 20 mg/ml collagenase (C8176, Sigma-Aldrich) in 0.05% trypsin/0.53 mM EDTA/1X HBSS (25-051-CL, Corning) for 25-40 min in a thermomixer, homogenising by gently pipetting every 5 minutes. The dissociated cells were transferred to 1X HBSS/10 mM/0.25% BSA (A3059, Sigma-Aldrich) to stop the reaction, rinsing the original tube to collect any residual cells. After homogenisation, cells were centrifuged at 500 g for 10 min at room temperature. Cells were resuspended in HBSS, filtered through a 40µm cell strainer (352340, Corning), centrifuged at 750 g for 10 min and FAC-sorted for GFP+ or mCherry+ (FACSAria, BD Biosciences Fusion System). Cells were counted before proceeding to single cell RNA sequencing.

### Single cell RNA library construction and sequencing

The Chromium Single Cell 3^*′*^ v3.1 platform (10X Genomics) was used for single cell RNA sequencing. Briefly, single cell suspensions for each condition were loaded onto the 10X Chromium Single Cell Controller using the NextGEM Chip G and the Chromium Next GEM Single Cell 3^*′*^ Reagent Kits v3.1 kits to capture single cells in individual droplets and obtain cell-barcoded mRNA. After reverse transcription, cDNA was amplified and libraries prepared and quantified using Qubit dsDNA HS Assay kit (ThermoFisher). Their quality was checked on a 2200 TapeStation system (Agilent D5000 ScreenTape System). All cDNA libraries were pooled and sequenced to a depth of 25,000 read pairs per cell using 150 bp paired-end sequencing on Illumina Novaseq 6000 by Source Bioscience (Cambridge, UK).

### scRNAseq data analysis

Sequencing data were processed using the 10X Genomics Cellranger (v7.1.0) pipeline. Fastq files were aligned to the Danio_rerio_GRCz11 UCSC reference genome using a custom reference compiled by the Cellranger mkref function. The reference was built from the Lawson zebrafish transcriptome annotation (60) and supplemented with EGFP and mCherry sequences incorporated into both the genome and annotation files. Following alignment and quantification, we recovered 2351, 2562 and 1616 high-quality cells from unopened, sham and cryoinjured samples, respectively. These samples showed median detection of 1711, 1590 and 2328 genes per cell, with corresponding median UMI counts of 6421, 5998, and 8872.

Matrices of each sample were first processed using CellBender to remove ambient and background RNA. Downstream quality control and preprocessing were performed in R (v4.4.3) using Seurat (v4.3.0.1) (61). Cells expressing fewer than 200 or more than 4,500 detected features, with more than 10% mitochondrial gene content, were excluded. Doublets were identified and removed using Scrublet (62). Data from each sample were normalised and variance-stabilised using SCTransform v2 (sctransform package v0.4.1) (63) and the resulting SCT-normalised matrices were used for sample integration. Principal component analysis (PCA) was performed on the integrated dataset and the first 30 principal components were used for UMAP visualisation, nearest-neighbour graph construction and Louvain clustering at a resolution of 1.0. All downstream analyses and visualisations in R/Seurat were performed using the SCT assay. Plotting was carried out using Seurat’s implementations, the scCustomize package (v1.1.3) (64), the SCP package (v0.5.6) (65), and ggplot2 for custom graphics.

Cell type annotation was performed using canonical markers for immune cell types including macrophages, B-cells and NK-like cells, as well as analysis of differentially expressed genes (DEGs) to uncover cell types. To identify the subpopulations of macrophages, clusters identified as macrophages were subset using the Subset function and differential gene expression across clusters was performed. Analysis of DEGs was conducted by investigating gene ontology enrichment, grouping of DEGs into functional groups and manual curation.

### Differential gene expression analysis

Differential gene expression (DGE) analysis was performed using Seurat’s FindMarkers and FindAllMarkers functions with the Poisson test, applied to the corrected expression matrices generated after PrepSCTFindMarkers function. DEGs were defined as genes with adjusted P < 0.05 and log2 fold-change 1.

### Gene ontology enrichment analysis

Gene Ontology (GO) enrichment analysis of differentially expressed genes (DEGs) was conducted using clusterProfiler (66) in combination with org.Dr.eg.db to map gene symbols. DEGs were defined as genes with adjusted P < 0.05 and log2 fold-change 1, comparing conditions as well as clusters depending on the analysis.

### Integration with published scRNAseq datasets

To incorporate structural cell types not represented in our dataset, we included a published single cell dataset of regenerating zebrafish hearts (40) (GSE159032), selected because its methodological approach and tissue source closely matched our studied samples. Briefly, in this study adult zebrafish hearts were collected from unopened, 3 days post cryoinjury and 7 days post cryoinjury conditions, as well as other timepoints. To provide broader coverage of non-macorphage cell types within the heart for downstream modelling, we integrated the unopened, 3 dpi and 7 dpi samples with our mpeg1.1+ derived datasets. Integration of datasets was performed using SCTransform v2, as described herein.

### Cell-cell communication analysis with NicheNet

To investigate ligand-receptor interactions, we used NicheNet (42) adapting the version 2 ligand-receptor-target gene databases for zebrafish by mapping human genes to their zebrafish orthologues via BioMart (67). Differential NicheNet analysis followed the developers’ differential NicheNet workflow, comparing regenerating and homeostatic conditions within each cell type or cluster for both sender and receiver populations. Niches were defined for each mpeg1.1^+^ population with more than 15 cells in the cryoinjury and unopened condition, where the structural cell types were set as senders and the mpeg1.1^+^ cluster was the receiver. The unopened and cryoinjury niches were then compared for differential expression and ligand-receptor-target gene inference. Differentially expressed genes between regenerating and homeostatic states were supplied to the ligand-receptor-target inference framework using the default Log2 fold change threshold of 0.15. All analyses were carried out on the SCT assay generated after SCTransform v2 normalisation.

### Visium spatial transcriptomics sample preparation and sequencing

Following extraction, injured and homeostatic hearts were kept in Ringer’s solution on ice until transfer to empty cryomolds, where excess liquid was removed and samples were embedded in fresh OCT. Each heart was oriented horizontally under a stereomicroscope before snap-freezing in liquid nitrogen-chilled isopentane, as per manufacture’s recommendations. Tissue blocks were stored at -80°C until further processing. For cryosectioning, tissue blocks and Visium Gene Expression slides (10X Genomics) were equilibrated in the cryostat at -20°C for 30 min prior to sectioning. Samples were mounted on retainers and sectioned at 10µm using an anti-roll plate. Sections from zebrafish hearts were placed onto the Visium Gene Expression slide within the designated fiducial frames. Sections were gently thaw-adhered to the slide surface before slides were transferred to 50 mL tubes and stored at -80°C until further processing.

Samples were processed according to the Visium Spatial Gene Expression User Guide (10X Genomics) using the Visium Spatial Gene Expression Kit (10X Genomics). Briefly, sections were fixed in chilled methanol for 30 min at -20°C, stained with hematoxylin and eosin, and mounted in 80% RNase-free glycerol for imaging. Imaging was performed on a Zeiss AxioScanner whole-slide scanner at 20X magnification. After imaging, sections were permeabilised at 37°C for 6 min. The ideal permeabilisation time was determined using the Visium Spatial Tissue Optimisation Kit (10X Genomics). After permeabilisation, the on-slide reverse transcription (RT) reaction was performed at 53°C for 45 min. Second strand synthesis was subsequently performed on-slide for 15 min at 65°C. All slide-based reactions were executed in a thermocycler fitted with a metal slide holder. Following second strand synthesis, samples were transferred to tubes for cDNA amplification and cleanup. Library quality control and quantification was assayed using TapeStation (Agilent) and Qubit (ThermoFisher) systems. 10X Genomics Visium libraries were pooled and diluted to a concentration of 6 nM in nuclease free water, followed by paired-end sequencing on an Illumina NovaSeq 6000 PE150 to a depth of approximately 57 million paired reads per sample. Sequencing parameters: Read 1: 28 cycles, i7 Index: 10 cycles, i5 Index: 10 cycles, Read 2: 90 cycles. The sequencing data was processed using the Space Ranger pipeline v1.0.0 (10X Genomics) using the Danio_rerio.GRCz11 reference genome with mCherry and GFP added and the ENSEMBL Danio_rerio.GRCz11.109 genome annotation.

### Spatial deconvolution of 10X Visium spots using Cell2Location

To spatially map annotated cell types from the integrated scRNA-seq dataset onto the Visium spatial transcriptomics (ST) data, we used cell2location v0.1.3. Briefly, Cell2Location (39) employs a Bayesian framework to estimate the spatial distribution of cell types by decomposing ST data into cell-type abundance profiles. Initially, cell2location uses negative binomial regression to infer cell-type-specific expression signatures from the reference scRNA-seq dataset. These signatures are then applied to the ST data for non-negative decomposition of mRNA counts at each spatial spot, providing spatially resolved estimates of cell type abundance. In addition to mapping, we applied the co-localisation functionality of cell2location to better understand the spatial organisation of cell types and predict potential cellular interactions. This analysis uses non-negative matrix factorisation (NMF) on the cell-type abundance estimates generated by cell2location. The NMF approach identifies groups of spatially co-localised cell types by decomposing the abundance profiles into additive components. These components captured patterns of co-localisation, reflecting the coexistence of multiple cell types and microenvironments at shared Visium locations.

### Merscope spatial transcriptomics gene panel design

Using the integrated single cell transcriptomics dataset consisting of mpeg1.1^+^ cells as well as reference structural cells, a 500 gene MERFISH panel was designed for high-resolution spatial transcriptomics using the Merscope platform (Vizgen). To capture the heart-specific cellular identities, bona fide markers that identify key structural populations were included. Genes with insufficient target regions and genes exceeding the abundance threshold based on our single cell RNA sequence data were removed from the panel in order to avoid optical crowding. Furthermore, key marker genes were identified to capture the heterogeneity of mpeg1.1^+^ cells identified from our single cell suspension data and be able to identify these cells *in situ*. For this, the presto package (68) was used to conduct differential gene expression analysis and a combination of Log2 Fold Change and area under the Receiver Operating Characteristic curve (auROC) (68) was used to select such marker genes. Predicted ligands, receptors and downstream target genes from our NicheNet analysis were curated using literature search and the top relevant genes were included in the panel design to capture potential niche-specific signalling interactions. Other biologically relevant genes involved in the post-injury response (reviewed in (3)) were also included. The probe panel was developed by Vizgen and its full description is included as Table S1.

### Fixed frozen Merscope heart sample preparation

Unopened and 5dpi cryoinjured wild type and csf1ra^j4e1/j4e1^ mutant hearts were collected, placed in ice-cold Ringer’s solution and transferred to 1X HBSS (14185052, Life Technologies) containing heparin (H3149, Sigma-Aldrich). Hearts were fixed in 4% PFA (043368.9M, Alfa Aesar) in 1X DPBS (14190144, Gibco) at 4ºC overnight and equilibrated in 15% sucrose (S9378, Sigma-Aldrich) for 6 hours at 4ºC under gentle agitation. Hearts were then transferred to 30% sucrose prior to embedding in OCT (361603E, VWR International) and frozen on dry ice. Tissue blocks were stored at -80ºC. Prior to sectioning, samples were equilibrated in the cryostat at -20ºC for 30 min. Serial 10µm sections were cut from each heart and mounted onto Merscope glass slides (Vizgen) pre-coated with 0.1mg/mL poly-L-lysine in RNase-free 1X DBPS. For RNA quality control, additional sections were cut and tested using the RNeasy Plus Micro Kit, Qiagen. Following sectioning, slides were left to dry in the cryostat chamber at -20°C for 30 min, then removed and equilibrated to room temperature. Sections were washed three times for 5 min in 1X DPBS, air-dried for 30 min and permeabilised in 70% ethanol at 4°C overnight in a parafilm-sealed Petri dish. Slides were subsequently stored in fresh 70% ethanol at 4°C, sealed with parafilm, until further processing. Merscope slides with sections were prepared following manufacture’s instructions (69). Briefly, heart sections were processed for DAPI and cell boundary staining, encoding probe hybridisation and post-hybridisation steps following Vizgen’s user guide (91600002 Rev F). A fresh gel-embedding solution was prepared for each run and heart tissues were cleared at 47°C for 1.5 h and then re-embedded in gel. This double-gel embedding minimised bubble formation. Slides were loaded into the Merscope flow chamber together with the corresponding gene-imaging cartridge for MERFISH acquisition.

### Segmentation of Merscope data

For segmentation, we used Baysor (v0.7.1) (70), a method that considers joint likelihood of transcriptional composition and cell morphology to segment based on transcripts detected. Each region within the Merscope data was segmented separately. After segmentation, outputted loom files were combined as Anndata objects and heart sections annotated for experimental condition and section identifier using the Python libraries plotly and dash (71, 72).

### Label transfer of cell type annotations for Merscope spatial transcriptomics

We used the single cell suspension transcriptomics dataset to annotate the spatial cells present in our MERFISH sections. The translating and cycling clusters were removed from the single cell suspension dataset. Since the macrophage compartment was included using our mpeg1.1^+^ cells, annotated macrophages from the reference samples were also excluded. We followed the integration and label transfer tutorial of scvi-tools (43). Briefly, single cell suspension (RNA counts) and spatial data were concatenated and cells with fewer than 10 counts removed. The data was normalised using Scanpy’s normalize_total function and subsequently log normalised using log1p. The single cell suspension RNA-seq samples and individual heart MERFISH sections were treated as batches for scVI integration. Subsequently, a scANVI model was trained and cell type annotation was predicted using this model.

### Transcript processing, analysis and visualisation of Merscope data

Detected transcripts were combined from all imaged regions and annotated for heart section and experimental condition. For this, the coordinate system of the segmented cells was used to draw bounding boxes and group detected transcripts. Subsequently, we developed plotting functions that use matplotlib (73). Plots were constructed in multiple layers, including background non-coloured genes and coloured genes.

Transcript quantification was performed by subsetting regions of interest using Plotly and Dash libraries. Within the subset transcripts, the ratio of gene of interest over total molecules captured was used per section, normalising for transcriptional load and as a proxy area of the selected region of interest. The ratio from multiple sections of the same condition was combined and standard deviation was calculated using Numpy.

### Spatial transcriptomics app for data visualisation in this study

We developed an interactive Plotly and Dash dashboard that provides access to the processed Merscope data from homeostatic and regenerating heart sections, including expression profiles for all 500 genes analysed in this study. This app is hosted on Hugging Face accessible at https://simoeslab-regenerating-heart-dashboard.hf.space/.

### Niche and microniche identification with NicheCompass

NicheCompass (v0.3.0) (46) was used to identify interacting niches from MERFISH spatial transcriptomics data. The graph deep-learning model encoded a latent embedding with heart sections as a categorical covariate. Default settings were employed for model training with exception to the edge batch size (2048). Whole heart niche identification used data integrated across experimental conditions, including homeostatic/unopened and regenerating hearts of both wild type and csf1ra mutant phenotypes. Latent embeddings were clustered by Leiden algorithm at a resolution of 0.5 then annotated based on cellular composition, anatomical region and experimental condition. Injury zone microniche identification used data from one section each of wild type and mutant regenerating hearts. Clustering was conducted at a resolution of 0.3. Enriched Gene Programmes (GPs) within microniches were identified by differential testing of cell-cell communication strengths, where |log K| ≥ 2.3 defines enrichment. To investigate these enriched GPs, gene expression of ligands and receptors and GP score of each cell in the injury zone were visualised and investigated. Furthermore, the Scanpy function rank_genes_groups was used for differential expression testing across conditions within niches and microniches using normalised, log1p counts and default settings. For visualisation of population-specific expression of ligands and receptors, raw counts of annotated cells were plotted using a vmax of p99 or 1 to maximise visibility. For visualisation of population-specific expression of ligands and receptors, raw counts of annotated cells were plotted using a vmax of p99 or 1 to maximise visibility.

As NicheCompass was initially programmed for human and mouse data analysis, we compiled a human-to-zebrafish ortholog dataset and incorporated package support for zebrafish data analysis. The ortholog dataset was retrieved from Ensembl BiomartServer and formatted to match NicheCompass input. NicheCompass GP dictionary building functions that retrieve prior knowledge of human databases (NicheNet and Omnipath) were adapted to incorporate our human-to-zebrafish ortholog dataset for ortholog mapping.

## DATA AVAILABILITY

Raw and processed single cell RNA sequencing data of mpeg1.1+ cells as well as the Visium Spatial Transcriptomics and processed Merscope data are available through the GEO Accession numbers GSE316360, GSE316465 and GSE316357.

## CODE AVAILABILITY

All code to process data and produce the results of this study is available on Github: https://github.com/simoeslab/cardiac_immune_niches.

An interactive app is published to explore the Merscope data and it is accessible through the following url: https://simoeslab-regenerating-heart-dashboard.hf.space/

## SUPPLEMENTARY DATA

The following supplementary tables are included in this manuscript:

Supplementary Information 1 - DEGs comparing conditions

Supplementary Information 2 - GO-terms comparing conditions

Supplementary Information 3 – DEGs per cluster

Supplementary Information 4 – Top NicheNet results

Table S1 - Merscope panel design

## ACKNOWLEDGMENTS

We would like to thank the Biomedical Services Unit for fish husbandry, the WIMM Flow Cytometry Facility, the MRC WIMM Advanced Single Cell OMICS Facility and the UCL UK Dementia Research Institute Single-Cell and Spatial Omics facility for excellent services. We thank Vizgen’s Field Application Scientist Dr Carole Chedid for technical expertise. This work was supported by BHF Research Fellowship FS/IBSRF/21/25088 to FCS, Oxford BHF Centre of Research Excellence studentship RE/18/3/34214 to ST and DPAG and Trinity College Whitehead Scholarship to ER.

## AUTHOR CONTRIBUTIONS

Conceptualisation, ER, ST, FCS; Methodology, ER, ST, TG, RA, FCS; Investigation, ER, ST, TG, FCS; Computational Analysis and Data Curation, ER, KC, WSW; Writing - Original Draft, FCS; Writing - Review Editing, ER, ST, TG, FCS; Supervision, RR, FCS; Funding Acquisition, FCS.

---

**Figure S1.**
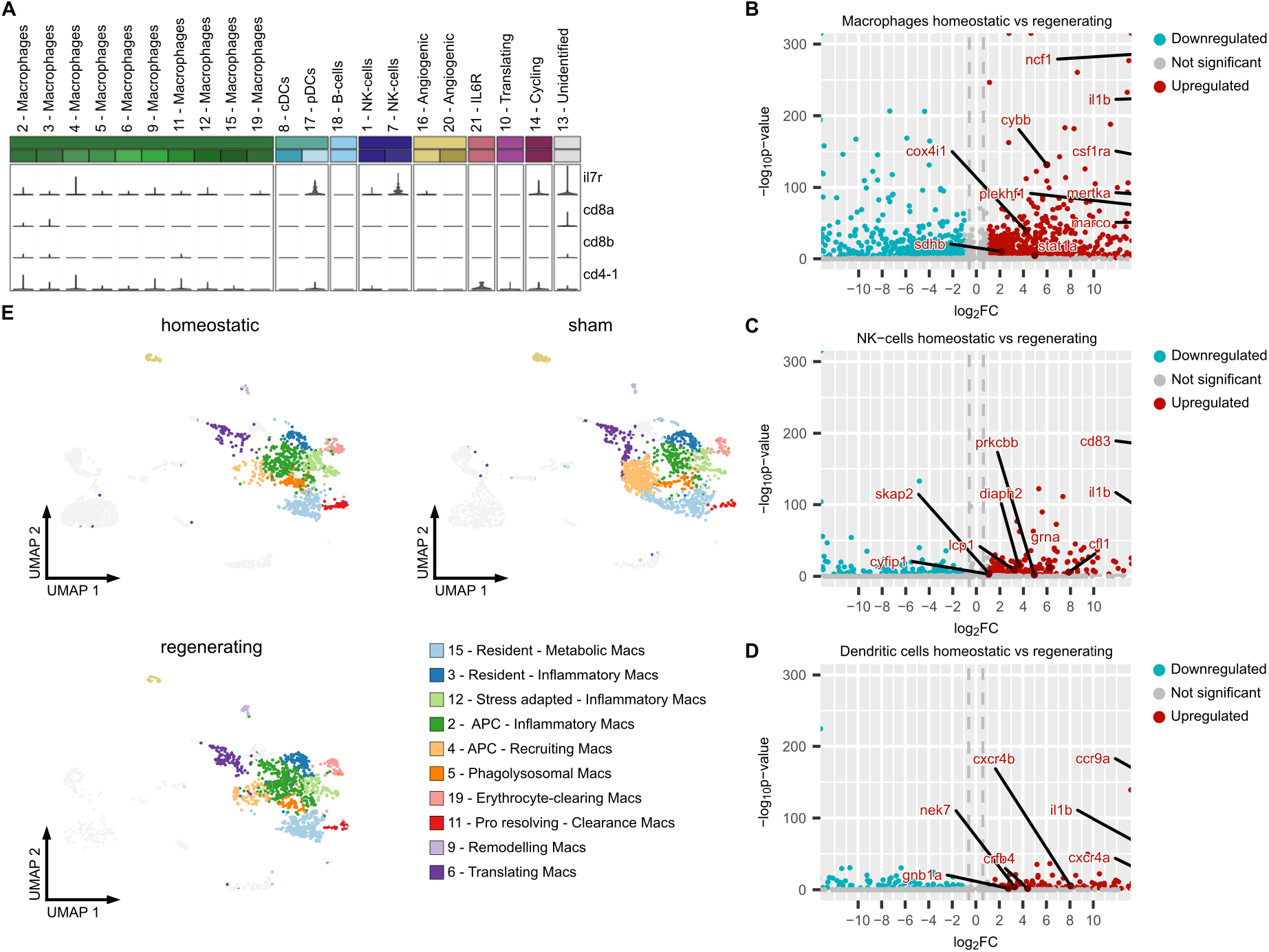
Additional characterisation of mpeg1.1^+^ populations and injury-induced transcriptional changes. A, Violin plots of T-cell markers (il7r, cd4-1, cd8a, cd8b) across mpeg1.1^+^ clusters, confirming no T-cell signatures were captured. B-D, Volcano plots of differentially expressed genes in homeostatic versus regenerating hearts for macrophages (B), NK-like cells (C) and DCs (D). E, UMAP embedding of macrophage subclusters coloured by condition (homeostatic, sham, regenerating), showing representation of each macrophage subpopulation across experimental groups.

**Figure S2.**
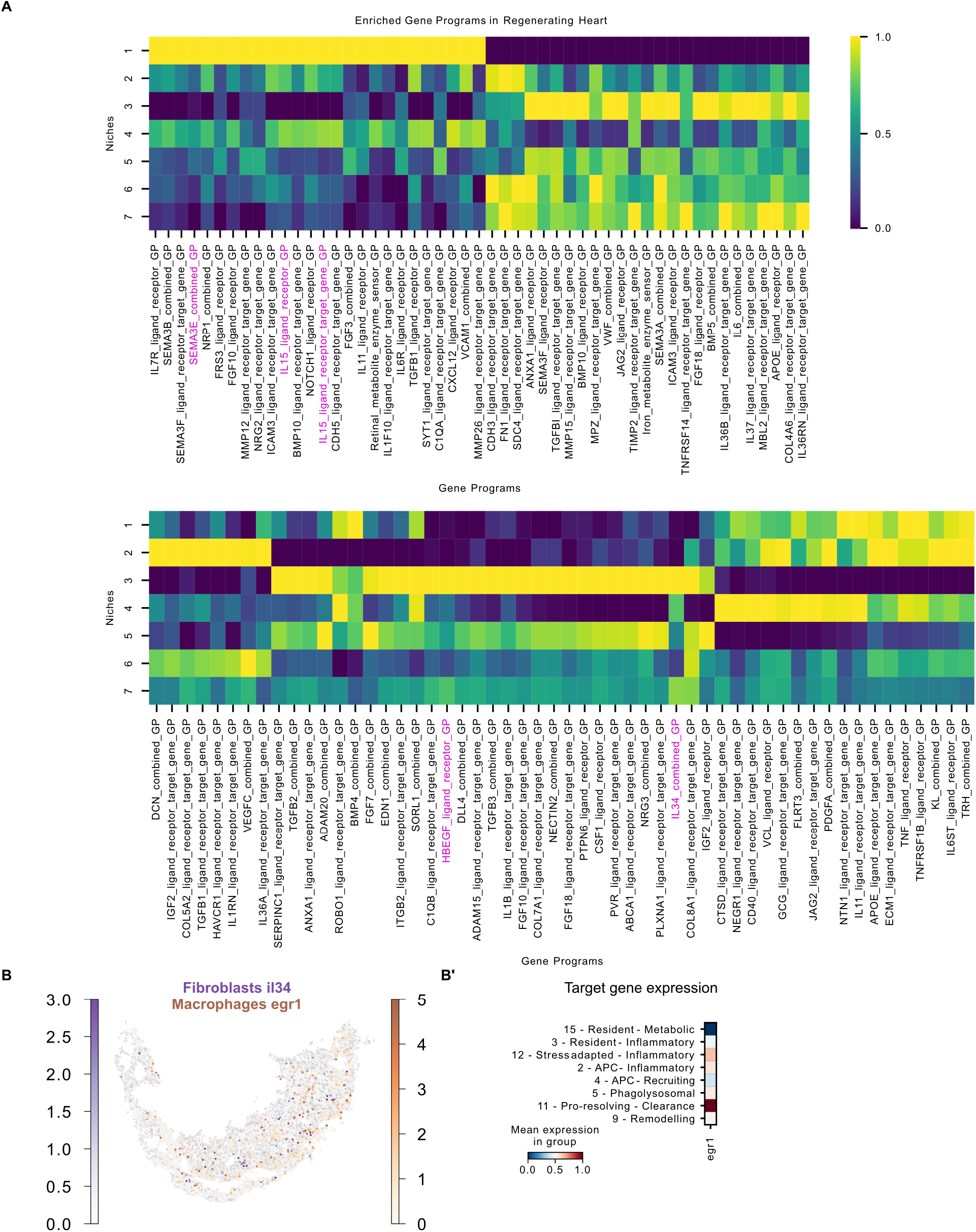
Gene programme landscape and il34-egr1 mapping across microniches. A, Heat map of NicheCompass-derived gene programmes enriched across microniches in the regenerating injury territory, highlighting microniche-specific combinations of signalling modules. B, MERFISH-based spatial co-localisation of il34 expression in fibroblasts and egr1 expression in macrophages across the injury zone. B*′*, Assignment of egr1 expression to macrophage subpopulations at single cell resolution.

**Figure S3.**
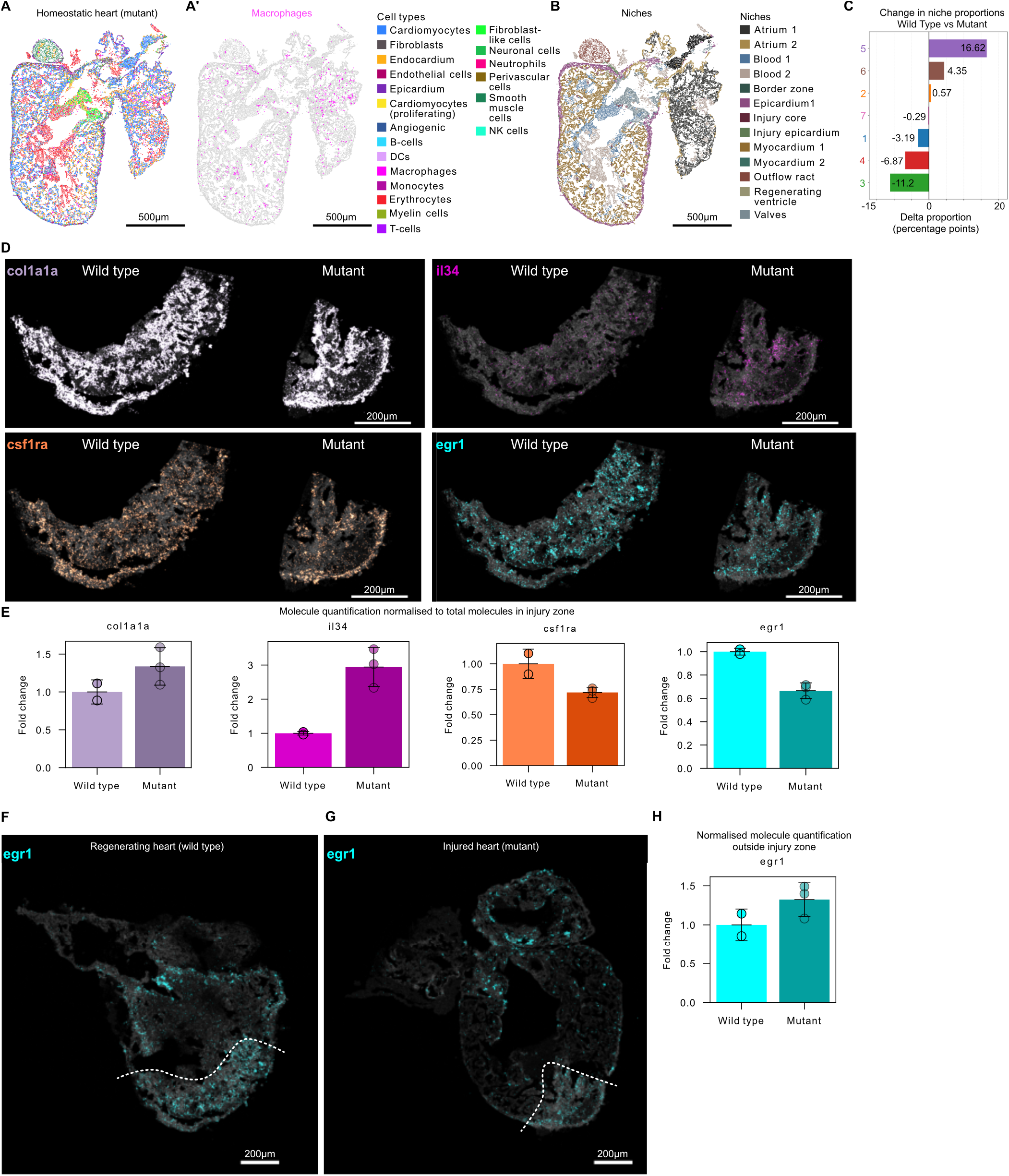
Steady-state niche architecture and il34-csf1ra-egr1 disruption in csf1ra mutants. A,A*′*, MERFISH map of the homeostatic csf1raj4e1/j4e1 heart coloured by cell type (A) and highlighting macrophages (A*′*). B, NicheCompass-derived niches in homeostatic csf1ra mutants, showing preserved global architecture. C, Change in microniche proportions within the injury territory of wild type versus csf1ra mutant. D,E, MERFISH spatial expression of col1a1a, il34, csf1ra and egr1 in wild type and csf1ra mutant hearts (D). Normalised fold change of captured col1a1a, il34, csf1ra and egr1 transcripts in mutant injury area compared to wild type. F,G, MERFISH whole-heart section of egr1 expression in wild type (F) and csf1ra mutant (G) hearts, showing that loss of egr1 is confined to the injury territory and not observed in remote myocardium. H, Normalised fold-change of egr1 transcripts in csf1ra mutants relative to wild type in regions outside the injury area.

